# A small RNA controls bacterial resistance to gentamicin during iron starvation

**DOI:** 10.1101/451765

**Authors:** Sylvia Chareyre, Frédéric Barras, Pierre Mandin

## Abstract

Phenotypic resistance describes a bacterial population that becomes transiently resistant to an antibiotic without requiring a genetic change. We here investigated the role of the small regulatory RNA (sRNA) RyhB, a key contributor to iron homeostasis, in the phenotypic resistance of *Escherichia coli* to various classes of antibiotics. We found that RyhB induces resistance to gentamicin, an aminoglycoside that targets the ribosome, when iron is scarce. RyhB induced resistance is due to the inhibition of respiratory complexes Nuo and Sdh activities. These complexes, which contain numerous Fe-S clusters, are crucial for generating a proton motive force (pmf) that allows gentamicin uptake. RyhB directly represses the expression of *nuo* and *sdh* operons by binding to their mRNAs, thereby inhibiting their translation. Indirectly, RyhB also inhibits the maturation of Nuo and Sdh by repressing synthesis of the Isc Fe-S biogenesis machinery. Notably, our study identifies *nuo* as a new direct RyhB target and shows that respiratory complexes activity levels are predictive of the bacterial sensitivity to gentamicin. Altogether, these results unveil a new role for RyhB in the adaptation to antibiotic stress, an unprecedented consequences of its role in iron starvation stress response.

**AUTHOR’S SUMMARY:** Understanding the mechanisms at work behind bacterial antibiotic resistance has become a major health issue in the face of the antibiotics crisis. Here, we show that RyhB, a bacterial small regulatory RNA, induces resistance of *Escherichia coli* to the antibiotic gentamicin when iron is scarce, an environmental situation prevalent during host-pathogen interactions. This resistance is due to RyhB repression of the synthesis and post-translational maturation of the respiratory complexes Nuo and Sdh. These complexes are crucial in producing the proton motive force that allows uptake of the antibiotics in the cell. Altogether, these data point out to a major role for RyhB in escaping antibacterial action.

## Introduction

The emergence and spread of bacterial multi-resistance to antibiotics has become a major health issue in the last decades, urging for the development of new anti-bacterial molecules and for a better understanding of the molecular mechanisms at work behind bacterial resistance (1, 2). While acquired resistance mechanisms (acquisition of genes or mutations that confer resistance) have long been the main focus of attention, less is known about “phenotypic” resistance, which is the process in which a bacterial population becomes transiently resistant to an antibiotic without requiring a genetic change (3-5). For instance, this kind of resistance has been associated with specific processes such as stationary growth phase, persistence and metabolic changes, reinforcing the idea that the environment encountered by the pathogen is a key determinant for antibiotic susceptibility (6).

Change in utilization of iron-sulfur (Fe-S) cluster biogenesis machineries in *Escherichia coli* gives a striking example of phenotypic resistance (7). Fe-S clusters are ubiquitous and ancient cofactors used in a plethora of biological processes, such as metabolism and respiration (8, 9). In *E. coli,* Fe-S clusters are formed and brought to target proteins thanks to two dedicated biogenesis systems: the so called “housekeeping” Isc machinery, which homologs are found in mitochondria of eukaryotic organisms, and the stress-responsive Suf system, in which homologs are found in chloroplasts of plants (10, 11). These systems are responsible for the maturation of more than 150 Fe-S cluster containing proteins in *E. coli,* notably numerous proteins contained in the main respiratory complexes I (Nuo) and II (Sdh) (12-14). Strikingly, it was shown that impairment of the *E. coli* Isc machinery enhances resistance to aminoglycosides, a well-known class of antibiotics that target the ribosome (7). This resistance is due to a deficiency in the maturation of the respiratory complexes in *isc* mutants, which in turn leads to a decrease in the proton motive force (pmf) that is essential for aminoglycosides uptake (15). Incidentally, it was deduced from these results that the Suf machinery is unable to maturate efficiently the Fe-S cluster containing proteins of the respiratory complexes, although the molecular reason for this still remains unclear. Overall this study predicted that an environmental signal that induces the switch from Isc to Suf should induce a transient resistance to aminoglycosides.

Iron starvation is one signal that decreases the expression of the isc operon encoding the Isc pathway. The small RNA RyhB mediates this regulation.(16). RyhB is one of the most studied sRNAs to date in *E. coli* (17-19). RyhB is regulated by Fur, the main regulator of Fe-homeostasis in many bacteria and is expressed during iron starvation (20, 21). When iron becomes limiting in the medium, RyhB base-pairs and represses the translation of more than 100 mRNA targets that encode for non-essential iron-utilizing proteins, thus engaging an “iron sparing” response and redirecting iron consumption in the cell (19). RyhB was shown to participate in the Isc to Suf transition during iron starvation by binding to the *iscRSUA* mRNA (16). In this way, it induces the degradation of the 3’ part of the mRNA that contains *iscSUA,* encoding the Isc machinery, while the 5’ part that encodes *iscR* remains stable. IscR is the major regulator of Fe-S clusters homeostasis and is itself a Fe-S cluster protein maturated by Isc (22). Accumulation of IscR in its apo-form has been shown to induce the *suf* operon (23). By its differential regulation of the *isc* operon, RyhB thus leads to the accumulation of apo-IscR that will turn on the expression of the alternative Suf system during iron starvation.

Iron homeostasis in particular has been shown to modify the sensitivity of bacteria to a number of antibiotics, although the molecular basis behind this is not always clear (24). Here we asked if the sRNA RyhB could participate in phenotypic resistance to various antibiotics during iron starvation. We found that RyhB is necessary to induce aminoglycoside resistance in low iron conditions. By further investigating the mechanism by which RyhB controls this phenotypic resistance, we show that RyhB controls entry of aminoglycosides in the cell by acting at both the synthesis and the maturation levels of the two pmf-producing respiratory complexes Nuo and Sdh.

## Results

### RyhB is involved in resistance to the aminoglycoside gentamicin

We first investigated the possible role of RyhB in resistance against different class of antibiotics during iron starvation. To do so, we performed antibiotic killing assays by growing wild type (WT) and *ryhB* mutant cells in LB medium starved or not for iron using 250 *μ*M of dipyridyl (DIP), a strong iron chelator. We chose this concentration of DIP because it is known to induce RyhB and we checked that it did not affect the growth of the cells (25, 26). Antibiotics were added when cells reached early exponential phase (OD_600_ = 0.2) and the number of survivors was determined by counting the number of colony forming units (c.f.u) after 3 hours of incubation. Four different major classes of antibiotics were tested: aminoglycosides (gentamicin), β-lactams (ampicillin), fluoroquinolones (norfloxacin), and tetracycline.

As expected, both WT and *ryhB* mutant cells were sensitive to the presence of all classes of antibiotics when grown in medium not starved for iron (Fig. 1A to D, left panels). Iron chelation did not protect cells against tetracycline (Fig. 1C). In contrast, adding DIP to the medium induced a protective effect on the WT and *ryhB* mutant strains for ampicillin and norfloxacin (Fig. 1A-B). The protective effect of iron deprivation for these antibiotics has already been observed and its underlying cause has been greatly debated (7, 24, 27, 28). As cells were protected independently of *ryhB,* we did not pursue these antibiotics further. In contrast, WT cells were protected against gentamicin when DIP was added to the medium, but this protection effect was lost when cells were mutated for *ryhB.* This result thus suggested that RyhB is involved in the protection of bacterial cells against aminoglycosides during iron starvation (Fig. 1D).

**Figure 1.**
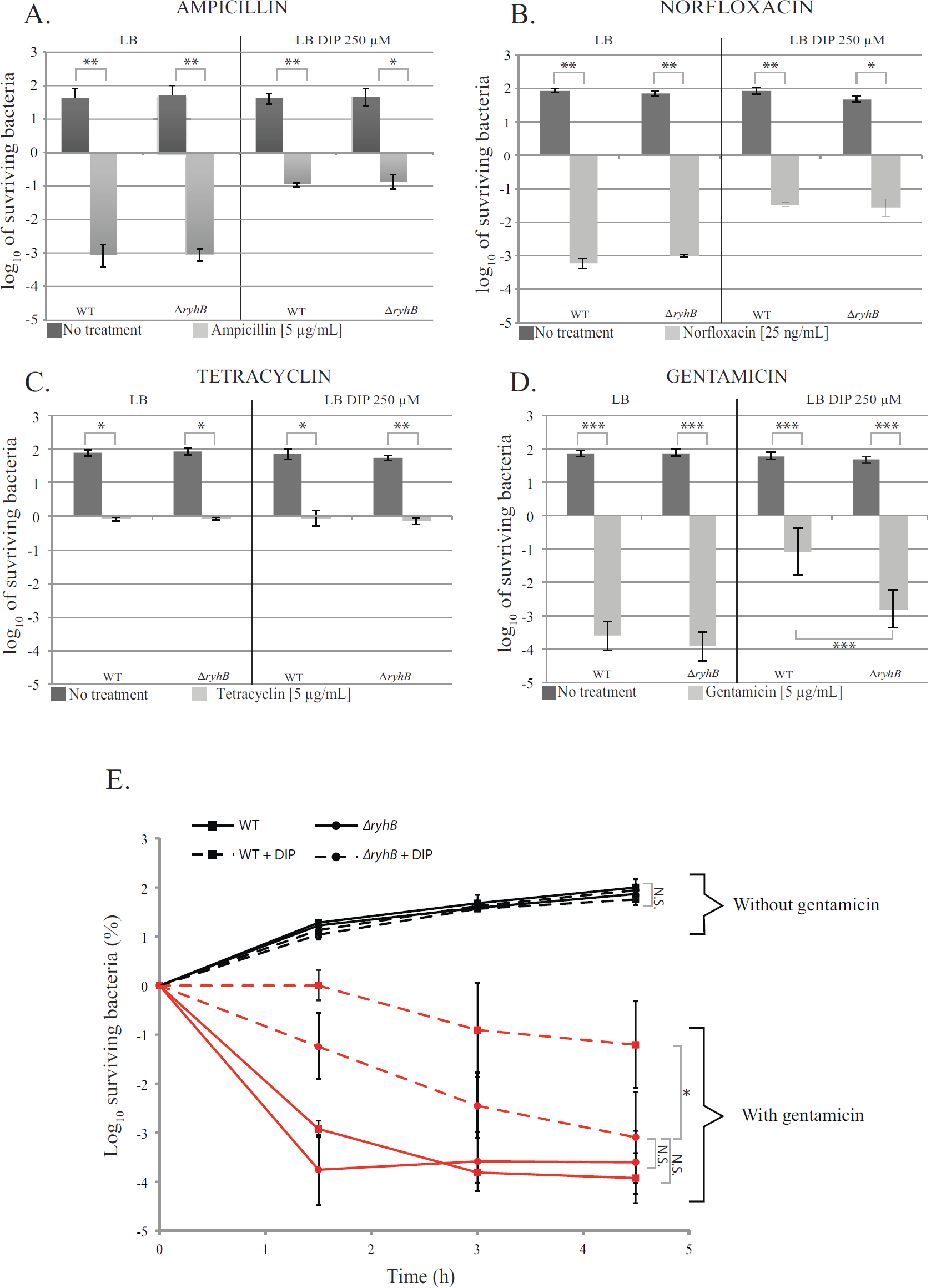
RyhB is involved in gentamicin resistance during iron starvation. A to D: strains were grown in LB (left panels) or in LB with DIP (250 *μ*M) (right panels) for 3 h with or without the following antibiotics A: ampicillin (5 *μ*g/mL); B: norfloxacin (25 ng / mL); C: tetracycline (5 *μ*g/mL) and D: gentamicin (5 *μ*g/mL). Colony forming units were counted to determine the number of surviving bacteria. Points were normalized relatively to to and plotted as log™ of surviving bacteria. The absolute c.f.u. at time-point zero was ≈ 5.10^7^ c.f.u. / mL for each sample. Error bars represent the standard deviations of three independent experiments. Statistical analysis were performed with Student’s T-test: *p < 0.05; **p < 0.01; ***p < 0.001. E: WT (squares) and *ryhB* mutant (circles) strains were grown in LB (regular lines) or LB depleted for iron (dashed lines) with (red curves) or without (black curves) gentamicin. The number of c.f.u. was determined at different times. Error bars represent the standard deviations of three independent experiments. Statistical analysis were performed with Student’s T—test: *p < 0,05; N.S.: Not significant.

To further investigate this phenotype, we performed gentamicin kinetic killing assays by growing WT or *ryhB* mutant cells in presence of DIP and counting the number of surviving bacteria at different time intervals after addition of the antibiotic. In this experiment, both the WT and *ΔryhB* strains showed the same profile when grown in LB (Fig. 1E). In both cases, the majority of the cells were rapidly killed after 1 h 30 min of incubation with gentamicin (5 logs of killing). Again, addition of DIP to the medium had a ≈ 4 log protective effect against gentamicin on WT cells as early as 1 h 30 min post addition of the antibiotic. Cells then remained mainly resistant to gentamicin during the course of the experiment. In contrast, the *ryhB* mutant gradually became as sensitive as cells grown in the absence of DIP (see 4 h 30 min time point), although killing kinetics were slightly slower than in presence of iron. Finally, to better characterize the effect of RyhB on gentamycin efficacy during iron starvation, we performed minimum inhibitory concentration assays (MIC) by growing WT and *ryhB* mutants in presence of increasing concentration of gentamicin, with or without DIP. Growing cells in presence of DIP almost doubled the MIC of the WT cells (from 6 *μ*g/mL in LB to 10 *μ*g/mL in presence of DIP) (Figure S1). In sharp contrast, the protective effect allowed by DIP was completely lost in the *ryhB* mutant. Altogether, these results indicated that RyhB is needed for the phenotypic resistance of *E. coli*to gentamicin in low iron condition.

### The RyhB induced resistance to gentamicin is dependent on Nuo and Sdh

Uptake of gentamicin has been shown to be a crucial step in the phenotypic resistance against this aminoglycoside (7). Entry of aminoglycosides is dependent on the proton motive force (pmf) mainly produced directly by respiratory complex I and indirectly by the respiratory complex II, respectively encoded by the *nuo* and *sdh* operon (12, 15, 29). Thus, one hypothesis was that RyhB induced resistance was due ! to an inhibitory effect on the activity of these two complexes that would block entry of gentamicin in the cell.

To test this hypothesis we repeated the previous killing assays in a strain deleted for both respiratory complexes *(Anuo Asdh).* As expected, this mutant was resistant to gentamicin (Fig. 2, left panel) (7). Adding DIP to the medium somewhat increased by 1 log the survival of the *nuo sdh* mutant, suggesting that pmf might be even more decreased in these conditions. Nevertheless, deleting *ryhB* from this strain did not increase its sensitivity to gentamicin during iron starvation (Fig. 2, right panel) indicating that the phenotype induced by RyhB was dependent on *nuo* and *sdh.*

**Figure 2.**
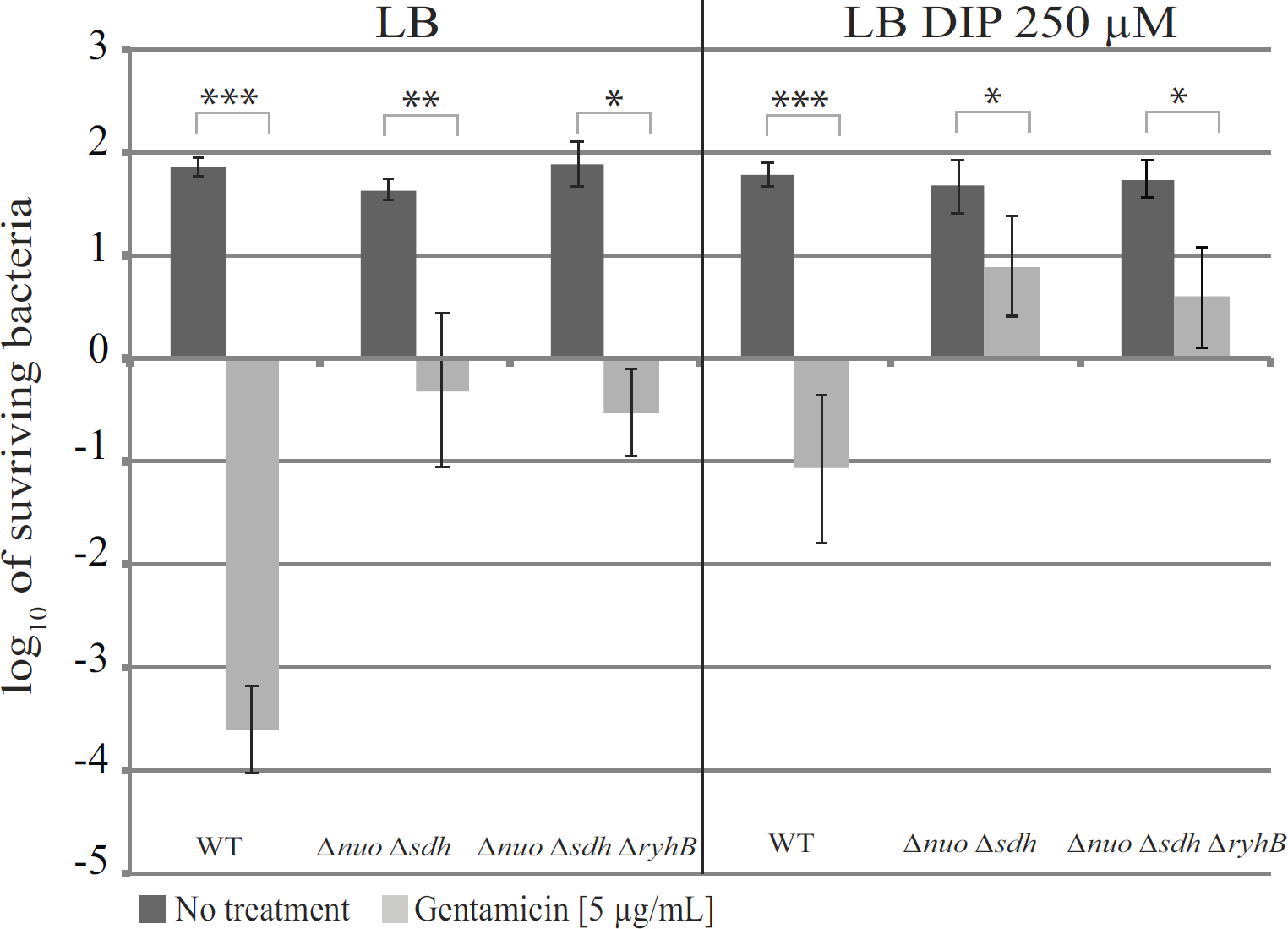
RyhB induced gentamicin resistance is dependent on Nuo and Sdh. The *Δnuo Δsdh* (BEFB20) and *Δnuo Δsdh ΔryhB* (SC024) strains were grown for 3 h with or without gentamicin (5 *μ*g / mL) in LB (left panels) or in LB with DIP 250 *μ*M (right panels). Colony forming units were counted to determine the number of surviving bacteria. Points were normalized relatively to t0 and plotted as log_10_ of surviving bacteria. The absolute c.f.u. at time-point zero was ≈ 5.10^7^ c.f.u. / mL for each sample. Error bars represent the standard deviations of three independent experiments. Statistical analysis were performed with Student’s T-test: *p < 0,05; **p < 0,01.

We further assessed the implication of each of the respiratory complexes by testing the sensitivity of the *Δnuo* and *Δsdh* simple mutants, deleted or not for *ryhB*(Fig. S2). The *nuo* simple mutant was almost completely resistant to gentamicin in presence of DIP, whether *ryhB* was present or not. In contrast, the *sdh* simple mutant became somewhat more sensitive (1 log) when *ryhB* was deleted from the chromosome. We conclude from these results that while both complexes are needed for full sensitivity of *ryhB* mutants to gentamicin, Nuo seems to be slightly more important than Sdh.

### RyhB represses the activity of the respiratory complexes

These previous results suggested that RyhB inhibits the activity of both respiratory complexes during iron starvation. To test this, we measured Nuo and Sdh specific enzymatic activities in WT and *ryhB* mutant strains grown in presence or absence of the iron chelator DIP in the growth medium. Nuo activity was decreased when the WT strain was grown in LB medium depleted for iron (about 4-fold) (Fig. 3A). In contrast, deleting *ryhB* from the chromosome restored 75% of Nuo activity in presence of DIP. The same pattern was also observed for Sdh activity (Fig. 3B). Altogether, these results confirm that RyhB represses the activities of both Nuo and Sdh complexes in medium deprived for iron.

**Figure 3.**
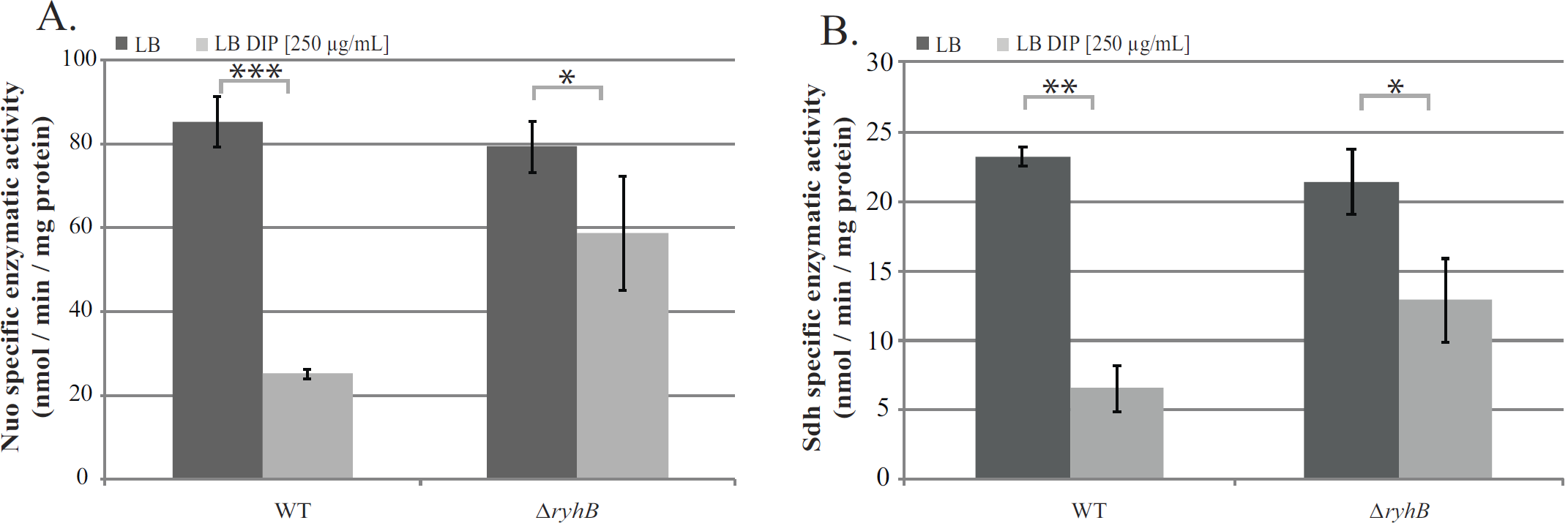
RyhB decreases Nuo and Sdh enzymatic activities. A: NADH specific enzymatic activity of Nuo in WT or *ΔryhB* strain grown in LB (dark grey bars) or in LB containing DIP (light grey bars) were determined by following the disappearance of the D-NADH substrate by spectrophotometry (nmol / min / mg protein). B: Succinate dehydrogenase activities in WT or *ΔryhB* strains grown in LB (dark grey bars) or in LB containing DIP (light grey bars) were determined by following the absorbance of DCPIP (nmol / min / mg protein). Bars represent the mean of at least three experiments and error bars represent the standard deviations. Statistical analysis were performed with Student’s T-test: *p < 0,05; **p < 0,01; ***p < 0,001.

### RyhB represses *nuo* and *sdh* expression

RyhB inhibition of Sdh and Nuo activities may be due to the repression of the synthesis and / or of the maturation of the complexes. Expression of *sdh* has already been shown to be repressed by RyhB (20, 30). In contrast, although pointed out in global approaches, RyhB regulation of *nuo* genes expression still awaited investigation (17, 31-33).

Using the RNA-fold software (http://unafold.rna.albany.edu), we could predict a base-pairing in between RyhB and the 5’ un-translated region of the first gene of operon, *nuoA* (34). This base-pairing involves 21 nucleotides (nt) of RyhB and includes the ribosome-binding site (RBS) and the start codon of *nuoA* (Fig. 4A). Overexpression of *ryhB* on a plasmid decreased the activity of a P_BAD_-*nuoA-lacZ* fusion of about 4-fold, as compared to cells transformed with an empty vector (Fig. 4B). In addition, the *P_BAD_*-nuoA-lacZ activity was decreased by 2-fold when WT cells were treated with DIP. This was in sharp contrast with the isogenic *ryhB* mutant strain for which activity remained the same in presence or absence of DIP (Fig. 4C).

**Figure 4.**
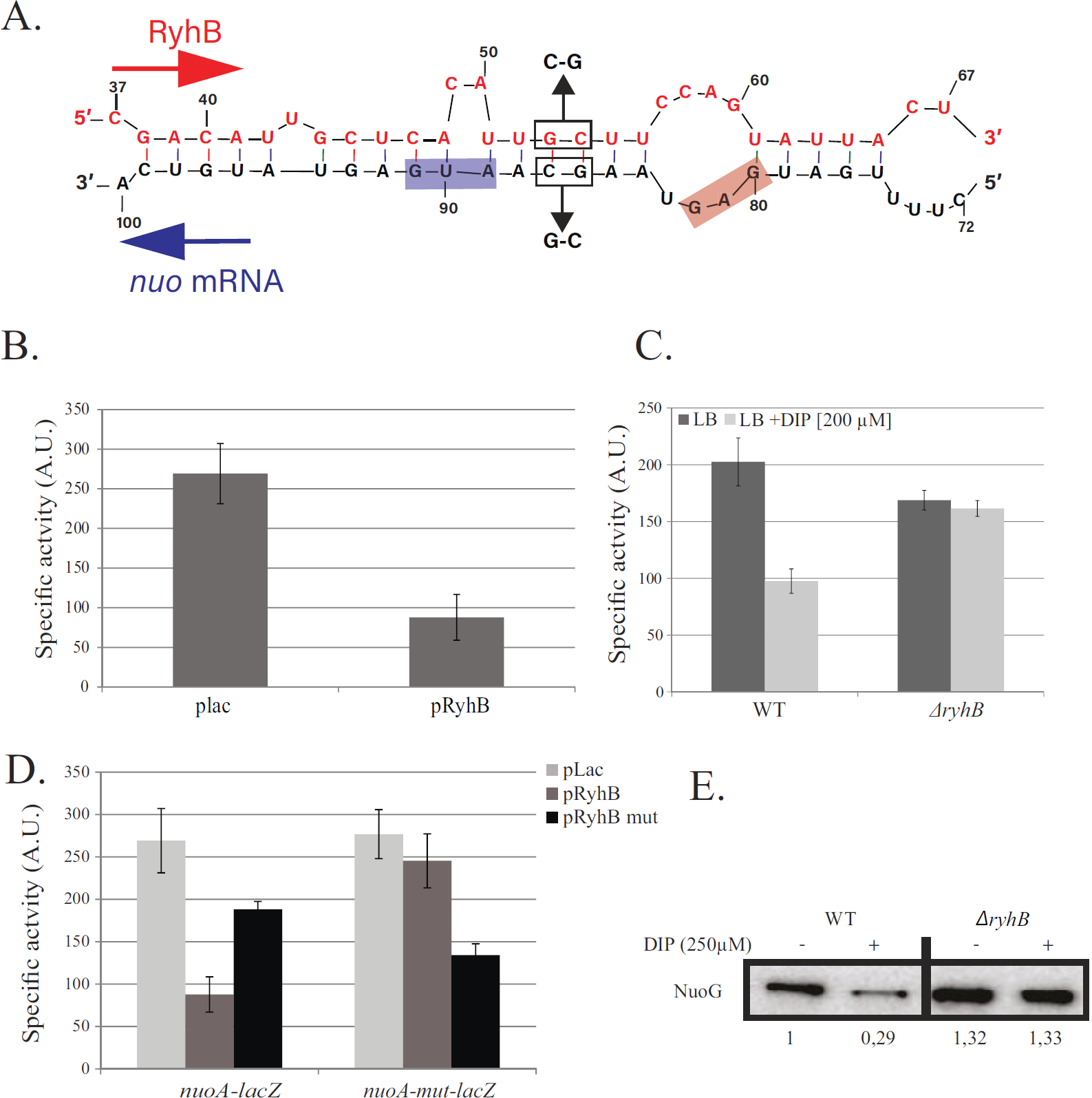
RyhB represses *nuo* expression. A: base-pairing predicted between RyhB and *nuo* mRNA. Nucleotides belonging to *ryhB* are represented on top, those corresponding to *nuo* on the bottom. Relative position to the transcriptional start site of *ryhB* and *nuo* are indicated above and below the sequences, respectively. B: the SC005 strain containing a P_BAD_-*nuoA-lacZ* fusion was transformed with the empty the plac vector or with the pRyhB plasmid containing *ryhB* under the control of an IPTG inducible promoter. Cells were grown in LB containing ampicillin (25 *μ*g/mL), IPTG (100 jUM) and arabinose (0,02 %) during 6 h after which 5-galactosidase activity was determined. Specific activities are represented by arbitrary units that were empirically determined to be approximately equivalent to Miller units. Error bars represent the standard deviations of six independent experiments. C: strains containing the P_BAD_-*nuoA-lacZ* fusion, WT (SC005) or deleted for *ryhB* (SC006) were grown in LB with or without DIP (200 *μ*M) during 6h before ß-galactosidase activities were measured. Each bar represents the mean from six independent experiments; error bars represent the standard deviations. D: Strains containing either the P_BAD_-*nuoA-lacZ* or the P_BAD_-*nuoA_mut_-lacZ* fusions were transformed with the plac, pRyhB or pRyhBmut plasmids and ß-galactosidase activity were determined. Each point represents the mean from six or more experiments. E: WT and *ryhB* mutant cell extracts from cultures grown in LB or in LB with DIP (250 *μ*M) were subjected to immunoblot analyses using antibodies raised against NuoG. Quantification represents the mean of three different experiments.

We then tested the biological relevance of the predicted base-pairing by introducing point mutations in the *P_BAD_*-nuoA-lacZ chromosomal fusion, giving rise to the *nuoA_mut_*-lacZ fusion (G86C and C87G; Fig. 4A). In contrast to the WT *nuoA-lacZ* fusion, RyhB overexpression was no longer able to repress activity of the *nuoA_mut_* fusion (Fig. 4D). We then introduced compensatory mutations in the pRyhB plasmid that should restore base-pairing to the mutated, but not to the WT, *nuo-lacZ* fusion, giving rise to *pRyhB_mut_*. As seen in figure 4D, overexpression of *RyhB_mut_* failed to fully repress the WT *nuo-lacZ* fusion, but was able to repress *nuoA_mut_-lacZ* fusion. Altogether these results show that RyhB represses *nuo* expression by base-pairing on the mRNA upstream *nuoA.*

We then evaluated the effect of this repression on protein levels by performing Western blot analyses against NuoG, a protein of the complex. Strikingly, NuoG protein levels decreased steeply, about 3-fold, when the WT strain was grown in presence of DIP (Fig. 4E). This phenotype was suppressed in the *ryhB* mutant, confirming the *in vivo* inhibition of Nuo synthesis by RyhB.

As a control and to compare *sdh* regulation to *nuo,* we performed a series of similar tests on an *sdhC-lacZ* fusion. We saw that RyhB overexpression repressed the expression of the fusion by more than 10 fold (Fig. S3A). In addition, the WT fusion was also strongly inhibited when cells were grown in presence of DIP but not when *ryhB* was deleted from the chromosome (Fig. S3B). Identical conclusions were reached from analyzing SdhB protein levels by performing Western blots (Fig. S3C). These experiments thus confirm the regulation of *sdh* by RyhB in our conditions.

### RyhB inhibits Nuo and Sdh maturation by repressing *iscSUA*

Biogenesis of Fe-S clusters by the Isc machinery has been shown to be key for full Nuo and Sdh activity and their associated pmf production. The *iscSUA* mRNA is a known RyhB target (16). Therefore, we asked if reducing levels of the Isc machinery synthesis following RyhB inhibition would be sufficiently important such as it would bear consequences on maturation of Nuo and Sdh.

To do so, we measured Nuo and Sdh specific activities in strains deleted for *suf* or for *isc* with or without *ryhB* (Fig. 5). In agreement with the literature, Nuo activity was decreased more than 5 fold in an *isc* mutant where the Suf machinery alone is responsible for Fe-S biogenesis (Fig. 5A). Activities of the *isc* mutant remained low in iron-deprived conditions, even when RyhB-mediated repression of Nuo and Sdh respiratory complexes synthesis was alleviated by deleting *ryhB.* Nuo activity of the *Δsuf* strain was comparable to that of the WT and DIP treatment inflicted the same drop in activity. Strikingly however, further deleting *ryhB* in the *suf*mutant almost completely restored Nuo activity when cells were grown in low iron condition. Thus, we concluded that repression of *iscSUA* and *nuo* by RyhB is sufficient to almost abolish Nuo activity. Moreover, these data strongly suggest that Isc is the only system that allows Fe-S clusters maturation of Nuo complex, regardless of the iron concentration in the medium.

**Figure 5.**
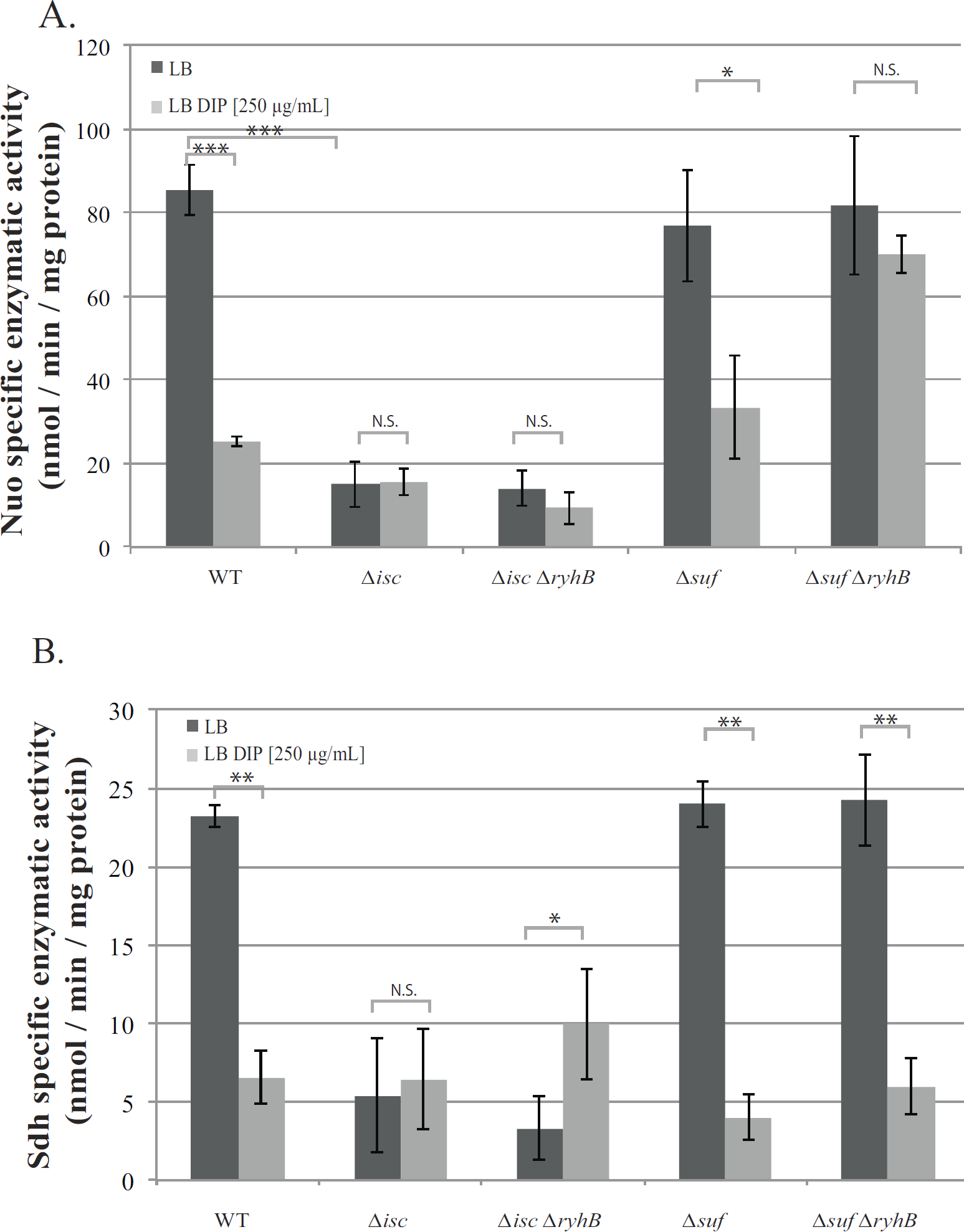
RyhB inhibits Nuo enzymatic activity by repressing *isc*. Nuo (A) and Sdh (B) specific enzymatic activities of *Δisc* and *Δsuf* mutants containing or not *ryhB* grown in LB (dark grey bars) or in LB containing DIP (light grey bars) were determined. Bars represent the mean of 3 independent experiments and error bars represent the standard deviations. Statistical analysis were performed with Student’s T-test: *p < 0,05; **p < 0,01; ***p < 0,001; N.S.: Not significant.

The situation was slightly different for Sdh. Deleting *isc* severely affected activity of Sdh in presence or absence of iron. In sharp contrast to Nuo however, activity of Sdh was not restored when *ryhB* was deleted from the *suf* mutant (Fig. 5B). Further deleting *ryhB* from this strain marginally restored Sdh activity, indicating that Suf can at least partially maturate Sdh proteins that are produced in absence of *ryhB.* These results thus suggest that Isc cannot ensure maturation of Sdh in low iron conditions.

### RyhB induces gentamicin resistance by repressing *isc, nuo* and *sdh* expression

In order to better appraise the role of Fe-S clusters maturation inhibition by RyhB in the resistance to gentamicin, we performed sensitivity assays in strains containing only one of the two Fe-S biogenesis machineries. As previously shown, the *isc* mutant was fully resistant to gentamicin in LB (Fig. 6A) (7). This phenotype remained unchanged when DIP was added to the medium, whether RyhB was present or not (Fig. 6A), thus showing that the slight Sdh activity observed in these conditions (Fig. 5) is not sufficient to render the cells sensitive to gentamicin. In sharp contrast, introducing a *ryhB* mutation restored sensitivity of a *suf* mutant strain when grown in presence of DIP (Fig. 6B), which is in agreement with the restoration of Nuo activity in this strain under these conditions.

**Figure 6.**
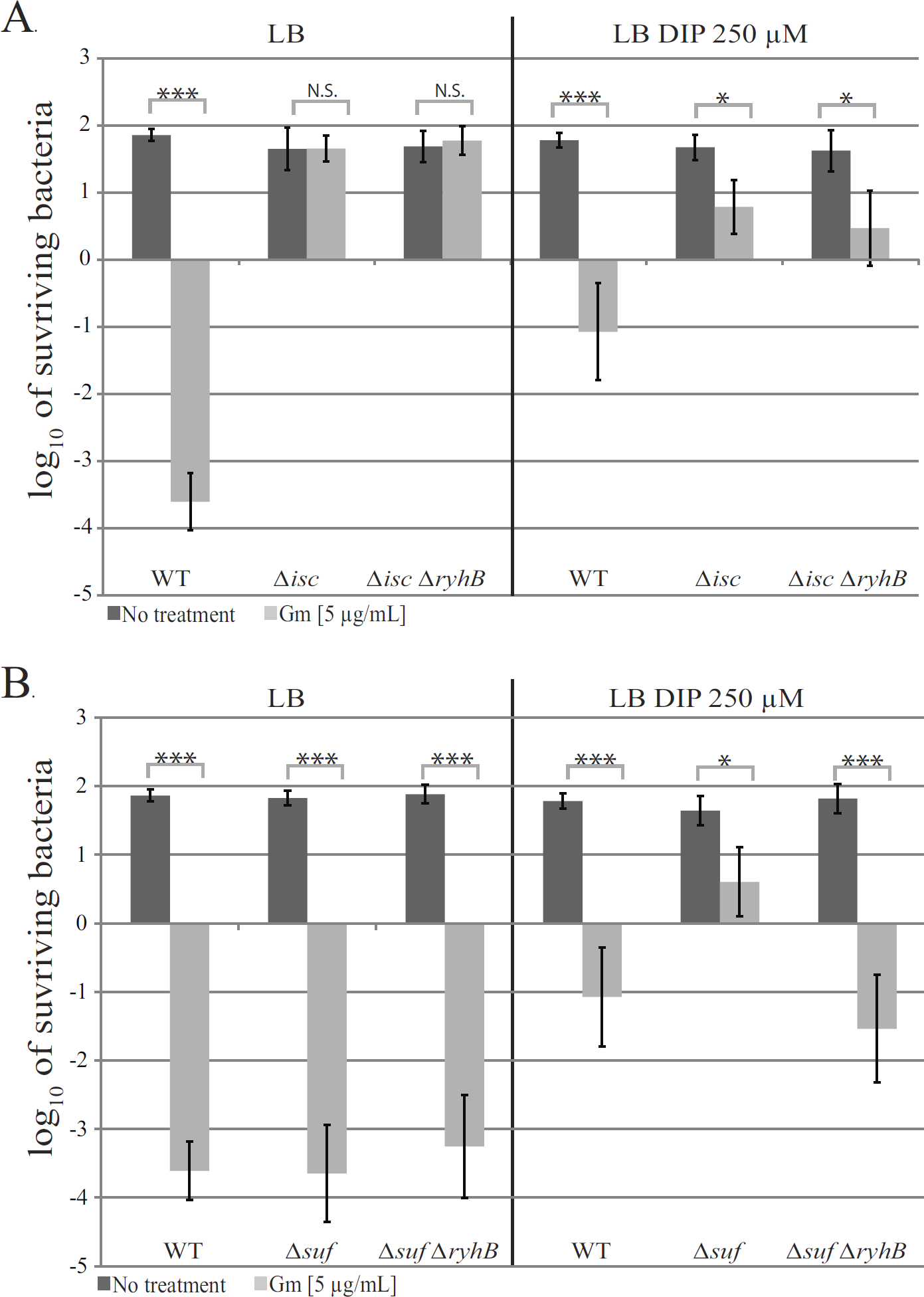
RyhB induces gentamicin resistance by inhibiting Fe-S clusters maturation. The *Δisc* (A) and the *Δsuf* (B) strains containing or not *ryhB* were grown with (light grey bar) or without (dark grey bars) gentamicin (5 *μ*g/mL) for 3 h in LB (left panels) or in LB with DIP (250 *μ*M) (right panels). After that, cells were diluted in PBS and spotted on LB agar plates. c.f.u. and Log_10_ of surviving bacteria numbers were determined. Error bars represent the standard deviation of three independent experiments. Statistical analysis were performed with Student’s T-test: *p < 0,05; ***p < 0,001; N.S. : Not significant.

As Nuo and Sdh activities are determinants for gentamicin sensitivity, we investigated if we could correlate both the levels of complexes enzymatic activity with that of resistance to gentamicin. Strikingly, there was an almost linear correlation between Nuo or Sdh activities of each strain and its sensitivity to gentamicin (Fig. S4 A and B). For instance, strains displaying the lowest Nuo activities were the most resistant to gentamicin, and vice versa.

Taken together, the ensemble of these results show that the maturation of Nuo and Sdh by Isc is essential for pmf production and that RyhB phenotypic resistance to gentamicin is due to both the direct inhibition of the expression of *nuo* and *sdh,* but also indirectly to the inhibition of Nuo maturation by Isc.

## Discussion

Phenotypic resistance can take place when environmental conditions change the metabolic state of the cell. Adaptative molecular responses modify cellular physiology, which induce a transient resistance state. Here, we show that the sRNA RyhB is a major contributor of *E. coli* phenotypic resistance to gentamicin in iron limiting conditions. Aminoglycosides uptake depends upon the activity of respiratory complexes I (Nuo) and II (Sdh) that produce pmf, directly and indirectly, respectively. RyhB acts negatively on both respiratory complexes, directly at the level of their synthesis and indirectly at the level of their maturation (i.e. acquisition of Fe-S clusters) (Fig. 7). Our model strengthens the role of the pmf-producing respiratory complexes in entry of aminoglycosides. Fe-S biogenesis maturation of the complexes was earlier pointed out as the main factor for resistance (7). By identifying here that the *nuo* mRNA is targeted by RyhB in addition to *sdh,* we show that synthesis of the respiratory complexes is also key in this process.

**Figure 7.**
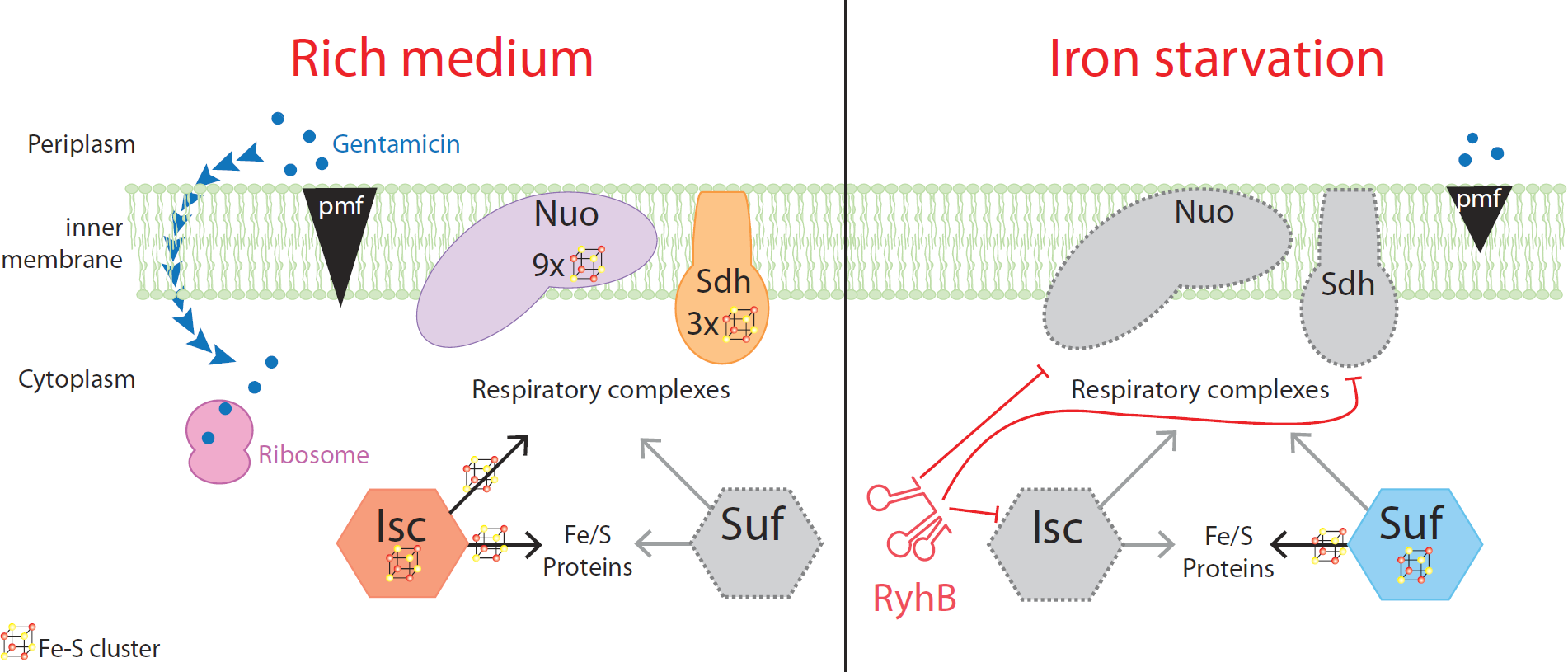
Model for the RyhB induced resistance to gentamicin during iron starvation. When iron is not limiting (left panel), the Isc Fe-S biogenesis machinery ensures the maturation of Nuo and Sdh, which generate a pmf that allows gentamicin uptake. Gentamicin reaches the ribosome and incudes mistranslation, which renders cells sensitive to the antibiotics. When iron is scarce (right panel), RyhB is expressed and represses the expression of *nuo, sdh* and *isc.* The pmf is lowered and gentamicin cannot enter the cytoplasm thus making cells resistant to the antibiotic.

As early as 2005, the *nuo* mRNA was suspected to be a target of RyhB as the operon was down-regulated when the sRNA was over-expressed, (17). The *nuo* mRNA was also more recently found associated with Hfq and RyhB in a global study of sRNA-mRNA interactions (33). We here could predict and confirm a direct base-pairing of RyhB to the *nuo* mRNA at the level of the UTR of *nuoA,* the first gene of the operon. This base-pairing occurs close to the ribosome binding site of *nuoA,* which strongly suggests that RyhB represses expression of *nuo* in a “classical” way, *i.e.* by occluding binding of the ribosome, leading to the degradation of the mRNA (35). The *nuo* mRNA is very long (about 15 kb) and comprises 14 genes, which makes it one of the longest mRNAs regulated by a sRNA to our knowledge. Importantly, in addition to the effects seen on *nuoA* by our beta-galactosidase assays (Fig. 4), we could also observe by Western blots RyhB repression on NuoG level (Fig. 4E), whose gene lies more than 5 kb away from the base-pairing site. It will thus be interesting to investigate how far downstream repression inhibits expression of the *nuo* operon.

Respiratory complexes are high iron consumers, with a total of 12 Fe-S clusters for Nuo and Sdh in *E. coli.* Thus, their repression by RyhB is in line with its role in installing an iron sparing response when iron becomes scarce (17, 19). Before our results, one could have imagined that RyhB represses Nuo and Sdh expression in order to limit accumulation of inactive apo-complexes in iron scarce conditions. However, both protein levels and activity of Nuo are restored in a *ryhB* mutant in iron-deprived medium indicating that maturation of respiratory complex I is possible under these conditions. These results strongly suggest that RyhB inhibits synthesis of Nuo Sdh not because they cannot be matured, but rather to preclude respiratory complexes to divert iron from other essential processes.

By repressing the *iscSUA* mRNA expression, RyhB also inhibits indirectly the maturation of Nuo (Fig. 3A and Fig. 5A). In contrast, maturation of Sdh was only partially restored in the *ryhB* mutant in presence of DIP (Fig. 3B) and, perhaps more surprisingly, this activity did not seem to be dependent on Isc but rather on Suf (Fig. 5B). More investigation is needed to understand the molecular basis for the difference in between Isc and Suf substrates preference. In any case, our results also clearly show that Nuo activity is more important than that of Sdh in installing a phenotypic resistance to gentamicin (Fig. S2). This may relate to pmf production by Nuo and Sdh. Indeed, Nuo, but not Sdh, directly translocates 4 protons across the membrane, while both indirectly contribute to pmf production by passing electons to cytochrome oxydase (12, 36).

The inhibition of respiratory complexes activity suggests that RyhB controls a complete metabolic shift during iron starvation, likely from respiration to fermentation. Although much needs to be done to assess this hypothesis, our recent survey indicates that a significant number of genes encoding Fe—S dependent enzyme of the TCA cycle are under the negative control of RyhB (19). Whether their maturation is also under RyhB influence via its control of the Isc system is an exciting issue to address.

Our study puts RyhB on the focus among a growing number of sRNAs that have been directly or indirectly linked to antibiotic resistance (36-38). However, in most of these cases phenotypes were derived from overexpression of the sRNAs and not relevant to physiological conditions. For instance, 17 out of 26 *E. coli* sRNAs that were assessed in a systematic manner against a variety of antibacterial effectors were shown to affect sensitivity to antibiotics when overexpressed, but few showed any phenotype when mutated (39).

A most spectacular case is represented by the role RyhB could play in the bacterial persistence of uropathogenic *E. coli* to different classes of antibiotics, among which included gentamicin (40). Persistence is a phenomenon in which a fraction of the bacterial population enters a metabolically inactive state that enables it to survive exposure to bactericidal antibiotics (41). Interestingly, in this study it was proposed that *ryhB* mutants would induce less persister cells because they display increased ATP levels and altered NAD+ / NADH ratios. In the light of our results, we believe these effects are explained by the fact that *ryhB* mutants probably display higher levels of Nuo, Sdh and Isc and therefore are more metabolically active, but also more prone to uptake the antibiotic. It is noteworthy that these experiments were conducted in rich medium not devoid for iron, and after long treatment with antibiotics (four days), which may explain low induction of RyhB in only a small percentage of bacterial cells that would then be able to resist antibiotics treatment in a persister-like manner.

RyhB homologs and paralogs are found in multiple other bacterial species, which suggests that many bacteria outside of *E. coli* may share the resistance mechanism that we describe here (19, 42-44). In particular, other pathogenic bacteria such as *Yersinia, Shigella* or *Salmonella* possess not only RyhB homologs, but also the Isc and Suf system and rely on Nuo and Sdh for respiration on oxygen (45, 46). RyhB has also been implicated in promoting sensitivity to colicin IA, which is not an antibiotic in a narrow sense, but a bacteriocin secreted by other species to outcompete bacteria sharing the same niches (47). In addition, RyhB has been shown to be involved in the virulence of *Shigella dysenteriae* by repressing the major virulence regulator *virB,* and the sRNA may be associated with the virulence of *Yersinia pestis,* as the expression of its two RyhB homologs (RyhB1 and RyhB2) increases in the lung of infected mice (43, 48). Altogether, these data point out for a major role for RyhB in escaping antibacterial action.

## Materials and methods

### Strains and culture

All strains used in this study are derivatives of *E. coli* MG1655 and are listed in Table S1. Strains were grown in LB broth (Difco), containing various concentrations of 2,2’-dipyridyl (DIP) (Sigma) when stated. Transductions with P1 phage were used for moving marked mutation as described previously in (49). The plac and pRyhB plasmids used in this study are described and have been transformed as previously described in (50). All oligonucleotides used are listed in Table S2.

### Antibiotic sensitivity experiments

Starting from overnight cultures in LB, strains were diluted 1/100 time in fresh medium containing or not DIP and grown aerobically at 37 °C with shaking until OD_600_ ~ 0.2. At this point, antibiotics were added to the cells (gentamicin: 5 μg / mL; ampicillin: 5 μg / mL; tetracycline: 5 μg / mL and norfloxacin: 25 ng / mL). After 3 h cells were taken, diluted in PBS buffer and spotted on LB agar plates and incubated at 37 °C for 16 h. Cell survival was determined by counting the number of colony-forming units per mL (c.f.u. / mL). The absolute c.f.u at time-point 0 was of ≈ 5 × 10^7^ cells / mL in all experiments.

### Minimum inhibitory concentration (MIC) determination

The MIC were determined as previously described (51). Briefly, each antibiotic containing-well of a 96-well micro-titer plate was inoculated with 100 μL of a fresh LB bacterial inoculum of 2 × 10^5^ c.f.u / mL. The plate was incubated at 37°C for 18 h under aerobic conditions. OD_600_ for each well was then determined by measuring the absorbance on a Tecan infinite 200. MIC was defined as the lowest drug concentration that exhibited complete inhibition of microbial growth.

### Fusions construction

The P_BAD_-*nuoA-lacZ* and P_BAD_-*sdhC-lacZ* fusions were constructed and recombined in PM1205 strain, as previously described (25). Briefly, sequences corresponding to *nuo or sdh* genes starting from its +1 transcriptional start up to 30 nucleotides downstream of the ATG codon were amplified using oligonucleotides P_BAD_-nuoA-F or P_BAD_-sdhC-F, and lacZ-nuoA-R or lacZ-sdhC-R, respectively. PCR amplifications were carried out using the EconoTaq DNA polymerase from Lucigen. The purified PCR products were then electroporated into strain PM1205 for recombination at the *lacZ* site. Recombinants carrying the desired fusions (SC005 and SC009) were selected on LB plates devoid of NaCl and containing 5 % sucrose, 0,2 % arabinose and 40 μg / mL X-Gal (5-bromo-4-chloro-3-indolyl-D-galactopyranoside). Blue colonies were chosen, and the resulting fusions were sequenced using oligonucleotides lacI-F and Deep-lac.

Overlap PCR was used to introduce point mutation in the fusion. The two PCR products corresponding to the sequence upstream and downstream of the desired mutation were amplified by PCR with oligonucleotides nuoAmut-F and Deep-lac, and LacI-F and nuoAmut-R containing the desired mutation and using genomic DNA from the SC005 strain as a template. The two PCR products were then joined by an overlap PCR using oligonucleotides lacI-F and Deep-lac. The resulting PCR products were purified, electroporated in strain PM1205 and sequenced as described above.

For point mutations in the pRyhB plasmid, the pRyhB plasmid was first purified from a WT (dam+) *E. coli* strain, and then amplified by PCR with oligonucleotides RyhBmut-F and RyhBmut-R, containing the desired mutation. The native plasmid was eliminated from the resulting PCR product by Dpn1 enzyme digestion for 1 h at 37 °C. Plasmids containing the desired mutation were then purified and transformed in SC005 and SC0026 strains.

### β-galactosidase experiments

Overnight cultures of different strains were diluted 1/100 times in fresh medium in culture flasks containing ampicillin and IPTG (isopropyl β-D-Ithiogalactopyranoside) or DIP when indicated. After ≈ 7 hours of growth 100 μL of cultures were dispatched in 96 wells microtiter plates (triplicates for each conditions). Absorbance at 600 nm was measured in a microtiter plate reader (Tecan infinite 200 ®). Then, 50 μL of permeabilization buffer were added in each well (100 mM Tris HCl pH 7,8; 32 mM Na_2_HPO_4_; 8 mM EDTA; 40 mM Triton; H_2_O milli Q) and the microtiter plate was incubated for 10 minutes at room temperature. O-Nitrophenyl-β-D-galactopyranoside (ONPG) was added to the solution and appearance of its degradation product was immediately determined by measuring the absorbance at 420 nm on a Tecan infinite 200 during 30 minutes. The specific activities were calculated by measuring the Vmax of the OD_420_ appearance divided by the OD_600_. Values were then multiplied by 100000, a coefficient that was chosen empirically to approximate Miller units.

### Nuo and Sdh enzymatic activities

The Nuo and Sdh enzymatic activities were determined as previously described (52, 53). Briefly, overnight cultures of the strains of interest were diluted 1/100 times in fresh LB medium containing or not 250 μM of DIP and grown at 37 °C with shaking until they reached OD_600_ ≈ 0.6. Cultures were pelleted by centrifugation (11 000 G, 10 min at 4 °C) and washed in phosphate buffer (50 mM pH 7,5). Cells were then lysed at the French press and 100 *μ*L were immediately frozen in liquid nitrogen before determining Nuo activity. Nuo enzymatic activity was determined at 30 °C by monitoring the disappearance of the specific Deamino-NADH (DNADH) substrate at 340 nm every 5 s during 10 min at 30 °C in a spectrophotometer.

For Sdh activity determination, lysate samples from French press were pellet by centrifugation (11 000 G, 10 min at 4 °C) and the supernatant was used for membrane fraction preparation by ultracentrifugation at 45 000 G at 4 °C during two hours. Pellets were then resuspended in phosphate buffer and kept in liquid nitrogen for later Sdh activity measurements. The enzyme was first activated by incubation in 50 mM Tris-HCl (pH 7.5), 4 mM succinate, 1 mM KCN for 30 min at 30 °C. The enzymatic activity was measured in the membrane fraction by monitoring Phenazine EthoSulfate (PES)-coupled reduction of dichlorophenol indophenol (DCPIP) at 600 nm, in a reaction containing 50 mM Tris-HCl (pH 7.5), 4 mM succinate, 1 mM KCN, 400 μM PES and 50 μM DCPIP.

The specific activities were calculated by measuring the Vmax divided by the protein concentration in total extracts evaluated by absorbance at 280 nm.

### Quantification of Nuo and Sdh protein levels by Western blot analyses

Total extracts and membranes preparation prepared for Nuo and Sdh activities were used for quantification of Nuo and Sdh protein levels, respectively. Total protein levels were determined by measuring absorbance at 280 nm on a spectrophotometer. Same amount of total protein level were migrated on polyacrylamide gels Tris-gly Sodium Dodecyl Sulfate (Novex 4-20 % Tris-Glycine Mini Gels) then, transferred on nitrocellulose membrane using Pierce G2 Fast Blotter (25 V, 1,3 mA, 7 min). Protein level were detected by incubating the membrane with α-NuoG or α-SdhB (1/1000) antibodies from rabbit and then by an α-rabbit antibody (1/1000) coupled with Hrp peroxidase. Signals were detected by chemiluminescence with Pierce ECL Western blotting system on an ImageQuant LAS 4000 camera. Quantification of protein levels was determined by measuring the specific signal intensity of the bands corresponding to Nuo and Sdh proteins with the ImageJ software. Intensities were normalized using an unspecific band detected by the same antibody.

## ACKNOWLEDGMENTS

We would like to thank A. Battesti for precious help with strain constructions and A. Huguenot for critical guidance with enzymatic activities assays. P.M., F.B. and S.C. work was funded by the Centre National de la Recherche Scientifique (CNRS) and Aix Marseille Université (AMU). S.C. is a recipient of a Fondation pour la Recherche Médicale grant (FDT20170436820).

---

**Figure S1. RyhB increases the resistance to gentamicin during iron starvation.** The WT and the *ΔryhB* mutant MIC were determined by growing cells in medium containing various concentrations of gentamicin and the iron chelator DIP (250 μM). The MIC was defined as the lowest drug concentration that exhibited complete inhibition of microbial growth

**Figure S2. Sensitivity of *nuo* and *sdh* simple mutants to gentamicin.** *Δnuo*(BEFB05), *Δnuo ΔryhB* (SC085), *Δsdh* (BEFB06) and *Δsdh ΔryhB* (SC086) strains were grown with or without gentamicin (5 *μ*g / mL) for 3 h in LB with DIP 200 μM. Colony forming units were counted to determine the number of surviving bacteria. Points were normalized relatively to t0 and plotted as log_10_ of surviving bacteria. The absolute c.f.u. at time-point zero was ≈ 5.10^7^ c.f.u. / mL for each sample. Error bars represent the standard deviation of three independent experiments. Statistical analysis were performed with Student’s T-test: *p < 0,05; **p < 0,01; N.S. : Not significant.

**Figure S3. RyhB represses *sdh* expression.** A: strain containing a P_BAD_-*sdhC-lacZ* fusion (SC009) was transformed with the empty plac vector or with pRyhB plasmid containing *ryhB* under the control of an IPTG inducible promoter. Cells were grown in LB containing ampicillin (25 *μ*g/mL), IPTG (100 μM) and arabinose (0,02 %) during 6 h after which 8-galactosidase activity was determined. Specific activities are represented by arbitrary units that were empirically determined to approximate Miller units. Error bars represent the standard deviations of six independent experiments. B: strains containing P_BAD_-*sdhC-lacZ* WT (SC009) or deleted for *ryhB* (SC010) were grown in LB with or without DIP (200 μM) during 6h before β-galactosidase activities were measured. Each bar represents the mean from six independent experiments. C: WT and *ryhB* mutant cell extracts from cultures grown in LB or in LB with DIP (250 *μ*M) were subjected to Western blot analyses using antibodies raised against SdhB. Quantification represents the mean of three different experiments.

**Figure S4. Gentamicin sensitivity can be directly correlated with Nuo and Sdh specific activities.** Sensitivity to gentamicin of WT, *ΔryhB, Δisc, Δisc ΔryhB, Δsuf* and *Δsuf ΔryhB* strains grown in LB (black points) or in LB containing DIP (red points) were plotted relatively to their Nuo (A) or Sdh (B) enzymatic activity respectively. The mean line represents linear correlation between the gentamicin sensitivity and complexes activities A : R^2^ = 0,86593; B : R^2^ = 0,77648. Error bars represent the standard deviation of three independent experiments.

**Table S1. Strains and plasmids used in this study.**

**Table S2. Oligonucleotides used in this study.**

